# A replicating recombinant vesicular stomatitis virus model for dairy cattle H5N1 influenza virus glycoprotein evolution

**DOI:** 10.1101/2025.02.27.640582

**Authors:** Lindsey R. Robinson-McCarthy, Kylie E. Zirckel, Holly C. Simmons, Valerie Le Sage, Kevin R. McCarthy

## Abstract

A panzootic of highly pathogenic avian influenza (HPAI) H5N1 viruses from clade 2.3.4.4b has triggered a multistate outbreak in United States dairy cattle and an unknown number of human infections. HPAI viruses are handled in specialized biocontainment facilities. Ethical considerations limit certain experimental evolution experiments aimed at assessing viral resistance to potential therapeutics. We have developed a replicating recombinant vesicular stomatitis virus (rVSV) where we replaced its glycoprotein with the hemagglutinin (HA) and neuraminidase (NA) genes of a 2.3.4.4b H5N1 virus (rVSV-H5N1dc2024), which is free of these constraints. This virus grows to high titers and encodes a fluorescent reporter to track infection. We demonstrate the utility of rVSV-H5N1dc2024 in neutralization experiments, evaluating antibody escape and characterization of resistance mutations to NA inhibitors. rVSV-H5N1dc2024 or similar viruses may accelerate efforts to develop and evaluate interventions against this emerging threat to human and animal health.

**IMPORTANCE:** Highly pathogenic avian influenza H5 viruses have spread globally, established sustained transmission in mammals and caused human infections. Research on these viruses is restricted to high biocontainment laboratories. We report the characterization and utility of a surrogate, replicating virus that displays the two key influenza glycoproteins, hemagglutinin and neuraminidase, that can be safely handled in most research laboratories. This virus is amenable for the evaluation of antiviral antibodies and small molecule inhibitors and the evolution of viral resistance to these agents. This virus can enable a wider range of researchers to study H5 viruses of pandemic concern.

## INTRODUCTION

Beginning in 2020 a lineage of highly pathogenic avian influenza (HPAI), H5 clade 2.3.4.4b, spread beyond Southeast Asia and initiated a global panzootic on six continents (1). Unexpectedly, these viruses achieved sustained transmission in United States dairy cattle (2). At current, 967 herds in 16 states have been afflicted (3). At least 69 documented human infections have been reported (4), however serologic surveys suggest the case burden is far higher (5-7). Laboratory studies of HPAI require specialized containment facilities (8, 9). Ethical considerations limit efforts to understand how these viruses may evolve in the presence of potential therapeutics. These limitations may delay efforts to develop and evaluate interventions against these viruses for humans and animals.

Replicating recombinant vesicular stomatitis viruses (rVSVs) are handled at lower biosafety levels and can be modified to express foreign glycoproteins (8, 10-18). In the absence of the VSV glycoprotein (G), cell entry is dictated by the foreign glycoprotein(s). For instance, rVSV expressing the Ebola glycoprotein (GP) was used to identify its cellular receptor (19). Because VSV is largely apathogenic in humans, rVSVs are often vaccine candidates, including vaccines against H5 HPAI, and an approved ebolavirus Zaire vaccine (Ervebo) (14, 20-23). Unlike lentivirus or other viral pseudotype systems, rVSVs are capable of sustained replication and are amenable to evolution studies. rVSVs expressing SARS-CoV-2 spike were used to evaluate monoclonal and serum antibody escape, enabling the identification of therapeutic candidates and future antigenic variants (24, 25).

We have generated, characterized, and demonstrated the utility of a rVSV that expresses a dairy cattle H5 clade 2.3.4.4b hemagglutinin (HA) and neuraminidase (NA) (rVSV-H5N1dc2024). This virus grows to high titer, encodes a green fluorescent protein to track infection, has a monoclonal antibody (mAb) neutralization profile similar to a matched influenza isolate, and is inhibited by a small molecule targeting the NA catalytic site. We show that rVSV-H5N1dc2024 evolves resistance to mAbs in experimental evolution studies and is suitable for evaluating NA inhibitor resistance mutations. rVSV-H5N1dc2024and similar viruses can accelerate response efforts to the 2.3.4.4b panzootic by expanding the number of laboratories that can work with this virus.

## RESULTS

### Recombinant VSV expressing influenza glycoproteins as a surrogate for dairy cattle HPAI

To generate a replication competent virus to study the entry and evolution of highly pathogenic avian influenza virus at biosafety level (BSL)-2 we used the rVSV platform. We generated a molecular clone of VSV, rVSV-H5N1dc2024, in which we replaced the VSV glycoprotein with HA and NA from A/dairy cow/Texas/24-008749-001/2024, a dairy cattle derived H5 clade 2.3.4.4b virus (Figure 1A). This infectious clone encodes an eGFP reporter. rVSV-H5N1dc2024 grows to titers of 3.2×10^8^ plaque forming units (pfu)/ml, within one log of rVSV-eGFP encoding its own glycoprotein (Figure 1A). When sequencing the virus, we noted a mixed population in early passages, with some sequences containing a stop codon in the HA cytoplasmic tail (Figure 1B). We plaque purified and grew stocks of full-length (fl) and cytoplasmic tail truncated (Δct) viruses. The Δct virus grows to titers approximately 1 log higher than the fl version, similar to rVSV-eGFP.

**Figure 1.**
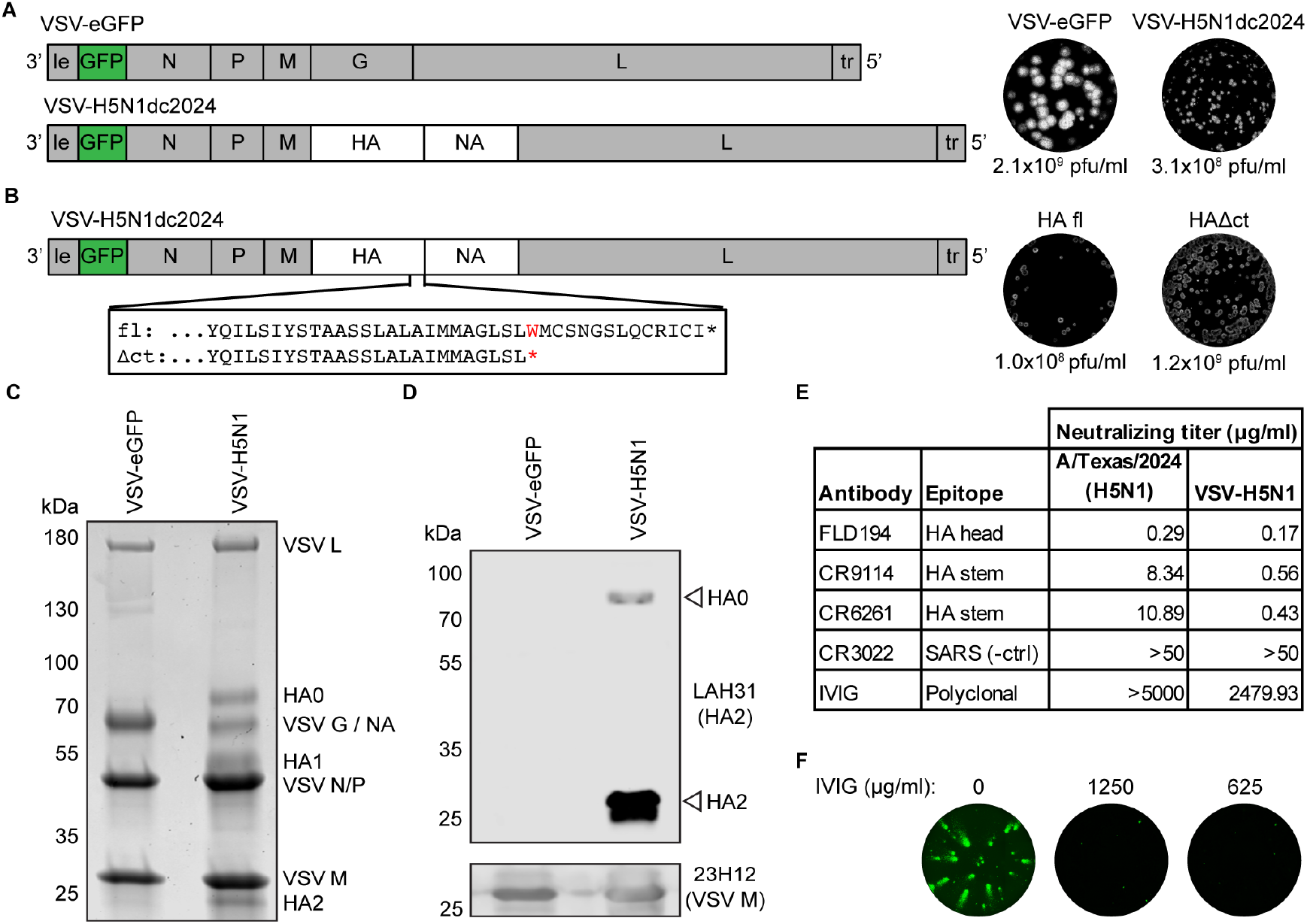
Generation of rVSV-H5N1dc2024. (A) Schematic of genomes, representative plaques, and infectious titers for rVSV-EGFP and rVSV-H5N1dc2024. (B) Identification of stop codon truncating the cytoplasmic tail of HA, representative plaques, and infectious titers of full length (fl) and cytoplasmic tail truncated (Δct) HA viruses. (C) Coomassie-stained SDS-PAGE of purified viruses. VSV G (only present in rVSV-EGP) and NA (only in rVSV-H5N1dc2024) run at the same position, as do VSV N and VSV P. (D) Western blot of HA incorporation into purified virions. Unprocessed HA0 and processed HA2 are present in viral particles. (E) Neutralizing activity of monoclonal antibodies and IVIG against authentic A/dairy cattle/Texas/24008749001/2024 (H5N1) and rVSV-H5N1dc2024. (F) IVIG inhibits spread of rVSV-H5N1dc2024. GFP-positive infected cells were imaged one day post-infection.

rVSV-H5N1dc2024 efficiently incorporates both HA and NA into particles (Figure 1C). In addition to the VSV core proteins M, N, P, and L, rVSV-H5N1dc2024 shows clear bands corresponding to HA and NA by SDS-PAGE. HPAI HAs are processed by furin proteases in the producer cell (26, 27). To verify that the bands we observed by SDS-PAGE correspond to unprocessed HA0 and processed HA1/HA2, we performed a western blot using antibody LAH31 which binds a conserved linear epitope in the long alpha helix of HA2 (28) (Figure 1D). The majority of H5 incorporated into rVSV-H5N1dc2024 particles is in its processed HA1/HA2 (fusion competent) form although some unprocessed HA0 is also present in virions.

We tested the susceptibility of rVSV-H5N1dc2024 to inhibition by H5-targeting mAbs and directly compared it to a sequence matched authentic A/dairy cow/Texas/24-008749-001/2024 virus generated through reverse genetics (29) (Figure 1E). In endpoint viral neutralization assays (29), rVSV-H5N1dc2024 and authentic H5N1 were inhibited by the same HA-specific mAbs and were not inhibited by SARS-CoV antibody CR3022 (30). rVSV-H5N1 showed similar sensitivity as authentic virus to mAbs targeting the HA head, and increased sensitivity to mAbs targeting the HA stem. Intravenous immunoglobulin (IVIG), purified IgG pooled from thousands of healthy human donors (31), did not inhibit authentic H5N1 and only had very weak activity against rVSV-H5N1dc2024. Although IVIG did not efficiently prevent infection, it did prevent spread of rVSV-H5N1dc2024 at sub-neutralizing concentrations (Figure 1F).

### Rapid generation of antibody escape mutants

rVSV-H5N1dc2024 is a replicating BSL-2 virus that cannot resort with circulating influenza viruses. Given these features, we demonstrate that this virus is suitable for assessing virus escape from mAbs. Using two potently neutralizing mAbs, which engage distinct but overlapping epitopes on the HA head (32, 33), we took two approaches to generate escape mutants (Figure 2A). First by “bulk” selection, pre-incubating virus with mAb, and growing the virus in the presence of the mAb. Second, by infecting cells, then adding mAb to the plaque assay overlay, and picking large plaques that formed (25). After four days of bulk selection with mAb FLD194, we observed viral spread throughout the culture (Figure 2B). Three days after plaque selection with both FLD194 and 65C6 we identified large plaques (4 and 1 respectively), which we picked and grew stocks from (Figure 2C). We identified three separate point mutations in FLD194 selected viruses (Q122R, Q122P, P125H). We identified K165M in the 65C6 selected virus. These mutations fall within the epitopes of the respective antibodies. (Figure 2D). All four mutant viruses were resistant to neutralization by the mAbs used in the selection experiment (Figure 2E).

**Figure 2.**
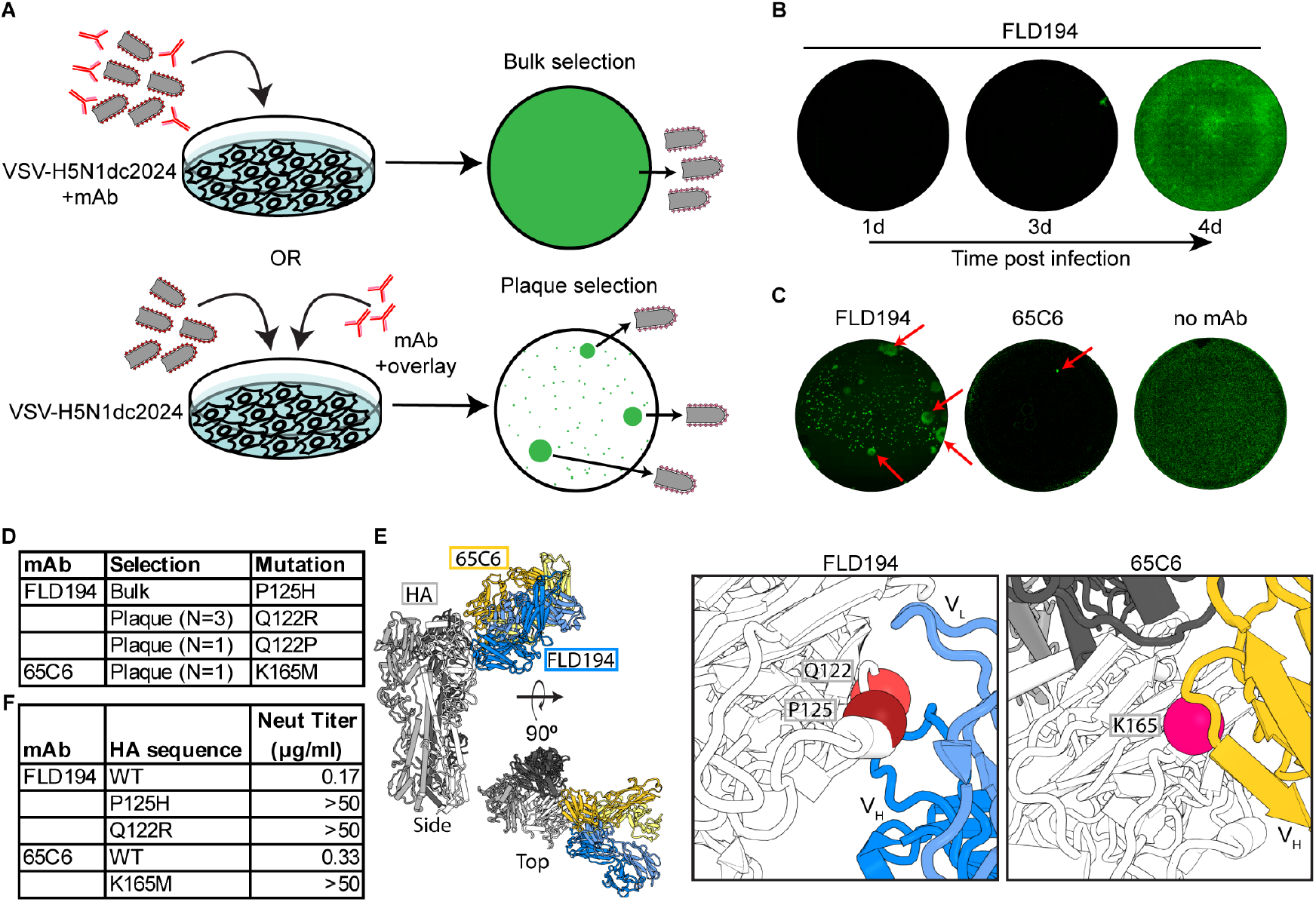
VSV-H5N1dc2024 for rapid selection of antibody escape. (A) Schematic of antibody escape selection strategies. (B) Bulk selection of resistance to FLD194. Cells were imaged over time to observe viral spread by 4 days post-infection. (C) Plaque selection of resistance to FLD194 and 65C6. GFP positive plaques were imaged 3 days post-infection. Arrows denote plaques that were chosen for further analysis. (D) Mutations identified in HA in selected viruses. (E) Structures of FLD194 and 65C6 complexed with an H5 HA (PDB: 5A3I, 9EKF). Left: superposition of FLD194 and 65C6 onto H5 HA, shown with both side and top views. Right: Position of mutations identified in mAb resistant viruses. Q122, P125, and K165 are shown as spheres. (F) Neutralizing titers of mutant viruses.

### Evaluating drug escape mutations

Neuraminidase inhibitors like oseltamivir inhibit virus release from producer cells. Replicating viruses are therefore the best tool to determine the direct effect of inhibitors of viral spread over multiple replicative cycles. Assessing the effect NA mutations identified by surveillance efforts by engineering these into the authentic virus requires extra scrutiny and biosafety/ethical considerations. NA H275Y confers strong resistance to oseltamivir, and was recently identified in clade 2.3.4.4.b viruses infecting domesticated poultry (34, 35). To determine whether H275Y confers resistance oseltamivir resistance in the context of clade 2.3.4.4b H5N1, we engineered this mutation into rVSV-H5N1dc2024 (Figure 3A). Oseltamivir reduces cell-to-cell spread of rVSV-H5N1dc2024 at concentrations of 2μM and above and does not inhibit rVSV-eGFP (Figure 3B). rVSV-H5N1dc2024-NA-H275Y is resistant to oseltamivir, exhibiting only partial inhibition at the highest concentration tested (500μM).

**Figure 3.**
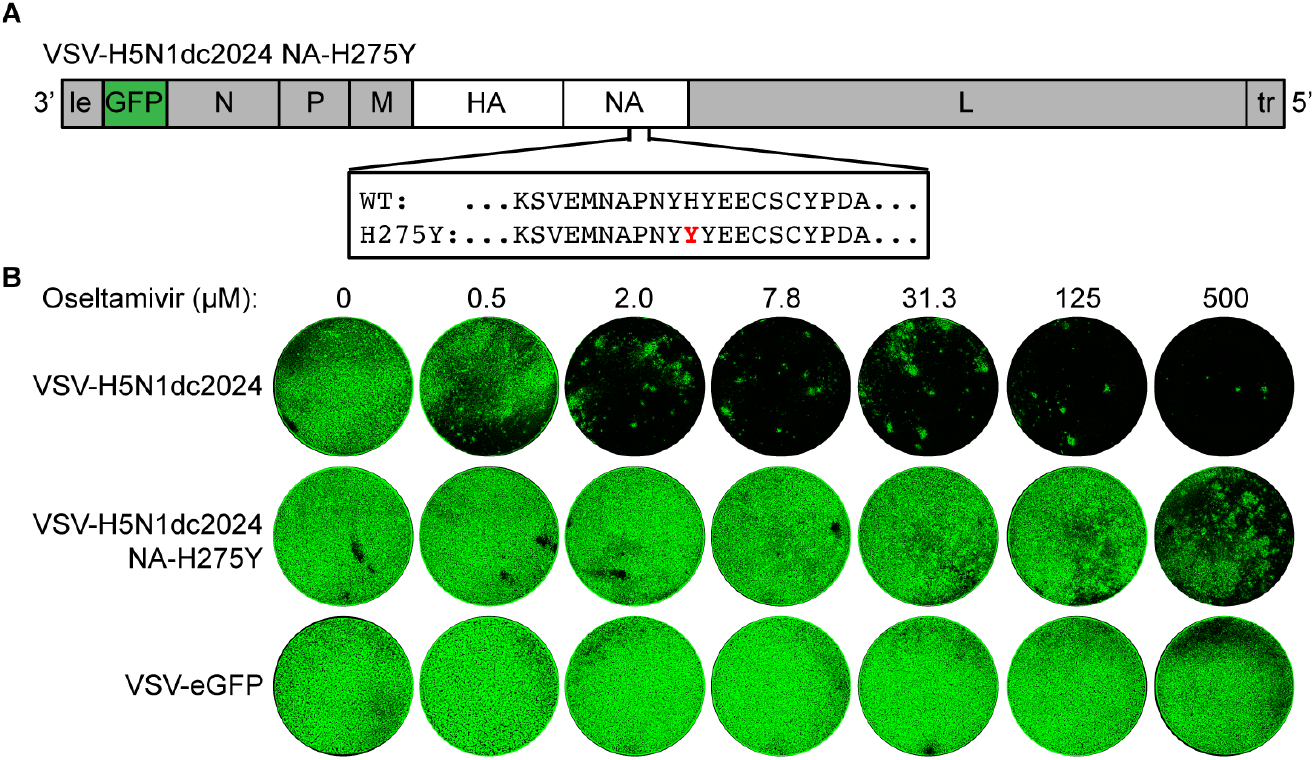
rVSV-H5N1dc2024 for assessing drug resistance. (A) Oseltamivir resistance mutation NA-H275Y was engineered into pVSV-H5N1dc2024-HAΔct. (B) Serial dilutions of oseltamivir were mixed with 100 pfu of the indicated virus. Infected cells were imaged for GFP 2 days post-infection to observe viral spread.

## DISCUSSION

The current outbreak of HPAI H5N1 continues to spread among dairy cattle, and the number of human cases of H5 clade 2.3.4.4b influenza infection continues to rise. The trajectory of this outbreak is unknown and experiments aimed at forecasting it are limited by requirements for high biocontainment and gain of function research concerns. To facilitate a rapid response to this potential pandemic pathogen, we produced a replicating BSL-2 virus for studying H5 clade 2.3.4.4.b HPAI glycoproteins. rVSV-H5N1dc2024 incorporates HA and NA into viral particles, grows to high titers, and has a similar pattern of neutralization by monoclonal antibodies as authentic matched influenza virus. We demonstrate that this system can be used to assess escape from monoclonal antibodies and evaluate resistance to antiviral drugs.

In humans, antibodies to HA are the major correlate of protection from influenza virus infection (36-38). rVSV-H5N1dc2024 has HA and NA sequences matching an authentic viral isolate. It recapitulates the neutralization phenotype of this authentic virus for mAbs targeting the HA head but is more sensitive to neutralization by mAbs targeting the HA stem. Further investigation is required to understand this cause of this discrepancy. Despite its heightened neutralization sensitivity to stem antibodies, we determined that human IVIG (produced from thousands of human donors) has little-to-no neutralization activity against rVSV-H5N1dc2024. The authentic virus was not neutralized. These observations agree with recent studies showing that individual donors have serum antibodies with limited-to-no neutralizing activity against H5 clade 2.3.4.4.b viruses (29, 39). Preexisting antibodies to this H5 are therefore likely to be lowly abundant in humans. Our use of a replicating virus with a fluorescent reporter enabled us to observe inhibition of cell-cell spread, suggesting the presence of antibodies in the human population that may act by interfering with viral assembly or egress.

Pandemic preparedness encompasses understanding the biology of an agent, developing therapeutics and vaccines, and evaluating the consequences of viral evolution in real time. We demonstrate that rVSV-H5N1dc2024, a BSL-2 agent, is a surrogate well-suited to rapidly prioritize experiments performed at higher biosafety levels. It is amenable to prospective studies of genetic barriers to therapeutic/prophylactic antibody resistance and to proactively assess known mutations that confer resistance to therapeutics. This includes assessing the effects of mutations, such as NA H275Y, which was recently identified in H5 clade 2.3.4.4b viruses in domesticated poultry (35). The virus itself is a potential vaccine candidate that can be manufactured at lower biocontainment and/or without the possibility of reassortment with circulating, seasonal, human, viruses (14, 18, 22). The malleability of rVSVs including H5N1dc2024 enable the rapid generation of panels viruses with mutations to rapidly assess their significance. The GFP reporter encoded by the virus is facilitates high throughput screening. Combined, rVSV-H5N1dc2024 and related viruses can accelerate pandemic preparedness and risk assessments.

## METHODS

### Cells

BSRT7 cells (40) and MDCK cells were maintained at 37°C and 5% CO_2_ in Dulbecco’s modified Eagle medium (DMEM; ThermoFisher) or minimal essential medium (MEM; ThermoFisher), respectively, supplemented with 10% fetal bovine serum (FBS) and penicillin/streptomycin (pen/strep; ThermoFisher). 293F cells were maintained at 37°C with 8% CO_2_ in FreeStyle 293 Expression Medium (ThermoFisher) supplemented with pen/strep.

### Plasmids

cDNA sequences corresponding to the HA and NA of A/dairy cow/Texas/24-008749-001/2024 was ordered from Integrated DNA Technologies (IDT). To generate pVSV-H5N1dc2024, HA and NA were cloned into pVSV-eGFPΔG (10) using MluI and NotI restriction sites. HA and NA were separated by the VSV intergenic sequence (TTTATGAAAAAAACTAACAGCAATC) and a KpnI restriction site. pVSV-H5N1dc2024-NA-H275Y was generated by site directed mutagenesis. The pVSV-H5N1dc2024-NA-H275Y plasmid contains a stop codon in the HA cytoplasmic tail to match the sequence of rVSV-H5N1dc2024-HAΔct. The pVSV-eGFP plasmid was previously generated (41). Sequences corresponding to the heavy and light chain variable domains of antibodies FLD194, CR9114, CR6261, CR3022, 65C6, and LAH31 (28, 30, 32, 33, 42, 43) were ordered from IDT and cloned into modified pVRC8400 plasmids containing full length human IgG1 heavy chains or human kappa or lambda light chains (44). All plasmid sequences were verified through Sanger sequencing (Azenta) or whole plasmid nanopore sequencing (Plasmidsaurus).

### Viruses

rVSVs were generated as previously described (45) with some modifications. BSRT7 cells were infected with Fowlpox-T7 (46) and transfected with VSV genomic plasmids along with helper plasmids encoding VSV N, P, L, and G. All rVSVs were propagated at 34°C on BSRT7 cells in DMEM supplemented with 2% FBS, 25mM HEPES, and pen/strep. Viral titers were determined by plaque assay in BSRT7 cells. Viral RNA was isolated using a QiaAmp viral RNA mini kit (Qiagen) and the region containing HA and NA reverse transcribed (Luna One-step RT-PCR kit, New England Biolabs). cDNA was sequenced using Oxford Nanopore technology (Plasmidsaurus).

A/dairy cattle/Texas/24008749001/2024 (H5N1) was previously generated through reverse genetics with sequences based on GISAID accession EPI_ISL_19014384 with noncoding regions determined from consensus alignment of H5N1 strains from the 2.3.4.4b clade viruses (29).

### Fluorescence Microscopy

rVSV plaque assays and infectivity assays were imaged for GFP expression using an EVOS automated fluorescence microscope (ThermoFisher) with a 4x objective. Images of whole wells were stitched together using integrated EVOS software and further processed using ImageJ software (National Institutes of Health).

### Monoclonal Antibody Production

IgGs were produced as previously described (47) by transient transfection of heavy and light chain plasmids into 293F cells using PEI transfection reagent. Five days post-transfection supernatants were collected, clarified by low-speed centrifugation, and incubated overnight with Protein A Agarose Resin (GoldBio) at 4°C. The resin was collected in a chromatography column and washed with one column volume of 10 mM tris(hydroxymethyl)aminomethane (tris), 150 mM NaCl at pH 7.5. IgGs were eluted in 0.1M Glycine (pH 2.5) which was immediately neutralized by 1M tris (pH 8.5). Antibodies were dialyzed against phosphate buffered saline (PBS) pH 7.4.

### SDS-PAGE and Western Blots

rVSVs were concentrated by ultracentrifugation over a 10% sucrose cushion and resuspended in PBS. Purified virus was boiled in Laemmeli buffer under reducing conditions and run on a 4-20% acrylamide gel (BioRad). Gels were stained with Coomassie protein stain and imaged with a LiCor Odyssey CLx imager (LICORbio). For western blot analysis, gels were run as described, then transferred to nitrocellulose, blocked in 5% nonfat dry milk in PBS with 0.1% Tween (PBST), and probed with mouse anti VSV-M antibody 23H12 (0.1 μg/ml; Milipore Sigma) and anti-HA2 antibody LAH31 (0.2 μg/ml) in 5% milk in PBST, followed by anti-mouse IR800 or anti-human IR800 (LICORbio) secondary antibodies. Membranes were imaged with a LiCOR Odyssey CLx imager.

### Neutralization Assays

A/dairy cattle/Texas/24008749001/2024 (H5N1) microneutralization assays were performed as previously described (29). Briefly, two-fold serial dilutions of monoclonal antibodies or IVIG (Gammagard 10% immune globubulin liquid, Takeda) were incubated with 10^3.3^ TCID_50_ of virus for 1 hour at room temperature with continuous rocking. Media was added to 96-well plates of confluent MDCK cells before the virus:antibody mixture was added. Cytopathic effect (CPE) was determined after four days, and neutralizing antibody titer was expressed as the reciprocal of the highest dilution of antibody required to completely neutralize infectivity. The concentration of antibody required to neutralize 100 TCID_50_ of virus was calculated based on the neutralizing titer dilution multiplied by the initial antibody concentration.

rVSV microneutralization assays were performed as above with modification. Two-fold serial dilutions of antibodies were incubated with 100 pfu of virus for 1 hour at room temperature with continuous rocking. Media was added to 96-well plates of confluent BSRT7 cells prior to addition of virus:antibody mixture. Infections was assessed by visual inspection for GFP positive infected cells after 1-2 days. Neutralizing concentration was determined as described above.

### Antibody Escape

For antibody escape in bulk, 10^6^ pfu of rVSV-H5N1dc2024 was incubated with 10μg/ml of FLD194 IgG for 1 hour at room temperature with continuous rocking. Media was removed from one well of a 6-well plate of confluent BSRT7 cells and replaced with virus:antibody mixture. Cells were incubated with virus:antibody mixture at 34°C and monitored for GFP expression every day until virus had spread throughout the culture at 4 days. Viral supernatant was collected and clarified with low-speed centrifugation and titer determined by plaque assay on BSRT7 cells. Viral genomic RNA was extracted and sequenced as described above.

For antibody escape by plaque selection, 6 well plates of confluent BSRT7 cells were infected with 10^6^ pfu of rVSV-H5N1dc2024 for 1 hour at 37°C. Virus was removed and agarose overlay containing 10μg/ml of FLD194 IgG or 65C6 IgG was added. Cells were incubated at 34°C and monitored for GFP expression. 3-5 days post-infection, large plaques were identified in each condition, picked, and grown on BSRT7 cells in the presence of 10μg/ml of the respective antibody. Viral stocks were titered and sequenced as described above.

### Oseltamivir Resistance Assays

Oseltamivir (Milipore Sigma) was reconstituted in sterile water. Serial 2-fold dilutions of oseltamivir in DMEM were mixed with 100 pfu of rVSV-H5N1dc2024, rVSV-H5N1dc2024-NA-H275Y, or rVSV-eGFP. Media was removed from 96-well plates of confluent BSRT7 cells and replaced with oseltamivir:virus mixture. Cells were imaged 2 days post infection and images processed as described above.

### Ethics and Biosafety

This project underwent evaluation by the University of Pittsburgh’s Dual Use Research of Concern (DURC) committee. It was determined that it was not a DURC concern and safe to perform at BSL-2. All experiments with authentic A/dairy cattle/Texas/24008749001/2024 (H5N1) virus were performed at BSL-3 at the University of Pittsburgh Regional Biocontainment Facility.

## ACKNOWLEDGEMENTS

We thank Sean P.J. Whelan and W. Paul Duprex for helpful discussions. This project was funded by funds from the University of Pittsburgh Center for Vaccine Research and by NIH award (UC7AI180311) from the National Institute of Allergy and Infectious Diseases (NIAID) supporting the Operations of The University of Pittsburgh Regional Biocontainment Laboratory (RBL) within the Center for Vaccine Research (CVR).

